## Supplementary main text for "Metabolic potential of uncultured bacteria and archaea associated with petroleum seepage in deep-sea sediments"

### List of contents

**Supplementary Note 1**

**Supplementary Note 2**

**Supplementary Table 1** Sample metadata and shotgun sequencing results.

**Supplementary Table 2** Alpha diversity estimates based on 16S rRNA gene amplicon sequencing.

**Supplementary Table 3** Relative abundance data for 16S rRNA gene amplicon sequencing and metagenome sequencing.

**Supplementary Table 4** Summary statistics for archaeal and bacterial MAGs.

**Supplementary Table 5** (1) GTDB-Tk classification of bins identified in this study as members of TA06; (2) Summary statistics of potential TA06 genomes based on NCBI and GTDB.

**Supplementary Table 6** Functional analysis of archaeal and bacterial MAGs. Presence/absence of genes are listed as: Presence: >1 (red), Absence: 0 (no color).

**Supplementary Table 7** Functional capacity based on normalized raw read count of genes (counts per million reads, CPM) encoding key metabolic enzymes.

**Supplementary Table 8** Accession numbers for genes involved in anaerobic hydrocarbon degradation used as custom database.

**Supplementary Figure 1** Map of the Eastern Gulf of Mexico (GoM) showing the studied three sampling locations (E26, E29 and E44) and bathymetry of the study area.

**Supplementary Figure 2** Phylum-level community composition of classified bacterial (left) and archaeal (right) 16S rRNA gene reads from separate bacterial and archaeal amplicon libraries. Detailed numbers can be found in Supplementary Table 3.

**Supplementary Figure 3** Phylogenetic relationship of putative glycyl-radical enzymes in MAGs with those of akyl-/arylalkylsuccinate synthases. Reference sequences show canonical alkane succinate synthase (AssA), 4-hydroxylbenzyl succinate synthase (HbsA), benzyl succinate synthase (BssA), and homologous putative alkane-degrading enzymes from *Vallitalea guaymasensis* L81 and *Archaeoglobus fulgidus* VC-16. The tree was constructed with the maximum-likelihood method (Poisson correction model) and is bootstrapped with 50 replicates. Sequences of pyruvate formate lyase (Pfl) from *E. coli* were used as an outgroup (not shown).

**Supplementary Figure 4** Phylogenetic relationship of identified genes in MAGs with currently-known molybdenum cofactor-containing hydrocarbon dehydrogenases. Reference sequences show the catalytic subunits of characterized cymene dehydrogenase (CmdA), alkane $C_2$-methylene hydroxylase (AhyA), and ethylbenzene dehydrogenase (EbdA/EbdA2) enzymes from other hydrocarbon-degrading bacteria. The tree was constructed with the maximum-likelihood method (Poisson correction model) and is bootstrapped with 50 replicates. Sequences of perchlorate reductase (PcrA) from *Dechloromonas aromatica* RCB were used as an outgroup (not shown).

**Supplementary Figure 5** Protein presence/absence matrix for benzoyl-CoA anaerobic biodegradation pathway. The MAGs were shown only if it was at least partially complete (presence of at least three subunits within one cluster for BcrABCD). Presence of genes is indicated by blue boxes. Gene names: Bcr, benzoyl-CoA reductase; Oah, 6-oxo-cyclohex-1-ene-carbonyl-CoA hydrolase; Dch, cyclohex-1,5-diencarbonyl-CoA hydratase; Had, 6-hydroxycyclohex-1-ene-1-carbonyl-CoA dehydrogenases. More details about these functional genes and pathways can be found in the text and in Supplementary Table 6.

**Supplementary Figure 6** Protein presence/absence matrix for reductive acetyl-CoA (Wood-Ljungdahl) pathway. Presence of genes is indicated by blue boxes.Columns correspond to the following enzymes: 1, formate dehydrogenase (Fhd) / formylmethanofuran dehydrogenase (Fwd); 2, formate-tetrahydrofolate synthetase (Fhs) / formylmethanofuran:tetrahydromethanopterin formyltransferase (Ftr); 3, methylene-tetrahydrofolate dehydrogenase (FolD) / N5,N10-methenyltetrahydromethanopterin cyclohydrolase (Mch); 4, methylene-tetrahydrofolate dehydrogenase (FolD) / methylenetetrahydromethanopterin dehydrogenase (Mtd); 5, methylenetetrahydrofolate reductase (Met) / N5,N10-methylenetetrahydromethanopterin reductase (Mer), 6, acetyl-CoA synthetase (Acs), 7 carbon monoxide dehydrogenase (Cdh).

**Supplementary Data 1** Genome sequences used for inferring Figure 2. It can be found at https://figshare.com/s/355963dc21a263e34c1f.

### Supplementary References

**Supplementary Note 1**

Among all identified phyla, candidate phylum TA06 is the only one not yet given provisional names. Also known as GN04 or AC1, it was originally discovered in a hypersaline microbial mat [1]. First genomic representatives of this phylum were recovered from estuarine sediments [2] with a small number of other MAGs recently reported to belong to this lineage [3, 4]. Due to the paucity of available MAGs and misclassifications based on 16S rRNA gene sequences, members of TA06 are often ‘confused’ with members of the phylum WOR-3 (*Stahlbacteria*) [4]. In addition to the phylogenetic inference here based on 43 concatenated protein marker genes (Figure 2), the placement of two bins within the original TA06 phylum is further supported by genome classification based on concatenation of 120 ubiquitous, single-copy marker genes [5] as well as classification of 16S rRNA genes using the SILVA database [6] (Supplementary Tables 4 and 5).

**Supplementary Note 2**

We performed a geochemical analysis of sediment porewater extracts. High concentrations of sulfate (Site E26: 16.55 mM; Site E29: 27.23 mM; Site E44: 25.63 mM) were detected at each of the three sites, consistent with sulfate being 28 mM in seawater and diffusing into sediments where it is consumed by sulfate reduction under anoxic conditions. $H_2$ and acetate concentrations were both below limits of detection (1 µM and 2.5 µM, respectively); this is consistent with previous observations in deep-sea sediments showing that $H_2$ and acetate are present at low steady-state concentrations due to tight coupling between producers and consumers [7, 8].

**Supplementary Table 1 Sample metadata and shotgun sequencing results.**

| ID | Site E26 | Site E29 | Site E44 |
|---|---|---|---|
| Latitude (N) | 26.59 | 27.43 | 26.28 |
| Longitude (W) | 87.51 | 86.01 | 86.81 |
| Geographic region | Henderson and Lund | Lloyd Ridge | Henderson and Lund |
| Water depth (km) | 2.8 | 3.2 | 3.0 |
| Quality-filtered reads | 85 825 930 | 148 908 270 | 138 795 692 |
| Contigs (> 500 bp) | 168 069 | 700 804 | 695 891 |
| Total size (bp) | 149 863 977 | 738 738 047 | 755 473 006 |
| Max contig (bp) | 52 576 | 191 288 | 346 360 |
| Average length (bp) | 891 | 1 054 | 1 085 |
| Curated MAGs | 9 | 37 | 36 |

**Supplementary Table 2 Alpha diversity estimates based on 16S rRNA amplicon sequencing.**

| | ***Richness Estimates*** | | | ***Diversity Indices*** | | | |
|---|---|---|---|---|---|---|---|
| **Locations** | **Observed** | **Chao1** | **ACE** | **Shannon** | **Simpson** | **Inversed Simpson** | **Fisher** |
| **16S amplicons of bacterial communities** | | | | | | | |
| Site E26 | 345 | 359 ± 7 | 356 ± 9 | 5.15 | 0.99 | 83.2 | 84.6 |
| Site E29 | 1055 | 1374 ± 41 | 1483 ± 21 | 5.97 | 0.99 | 135.9 | 412.4 |
| Site E44 | 348 | 360 ± 6 | 355 ± 9 | 5.28 | 0.99 | 100.8 | 85.6 |
| **16S amplicons of archaeal communities** | | | | | | | |
| Site E26 | 192 | 195 ± 3 | 196 ± 7 | 4.22 | 0.96 | 23.3 | 43.7 |
| Site E29 | 178 | 180 ± 2 | 181 ± 7 | 4.33 | 0.97 | 30.4 | 39.7 |
| Site E44 | 220 | 247 ± 12 | 241 ± 7 | 4.21 | 0.96 | 25.6 | 52.2 |

**Supplementary Table 7 Functional capacity of sediments based on normalized raw read count of genes (counts per million reads, CPM) encoding key metabolic enzymes.**

| ID | Site E26 | site E29 | Site E44 |
|---|---|---|---|
| ***assA-like*** | 233 | 394 | 346 |
| ***canonical assA*** | 9 | 37 | 10 |
| ***nmsA*** | 0 | 0 | 0 |
| ***bssA*** | 2 | 4 | 2 |
| ***acsB*** | 1485 | 2819 | 2474 |
| ***dsrA*** | 357 | 603 | 548 |
| ***FeFe*** | 332 | 380 | 335 |
| ***NiFe Group 1*** | 416 | 431 | 296 |
| ***NiFe Group 2*** | 55 | 5 | 23 |
| ***NiFe Group 3*** | 538 | 766 | 683 |
| ***NiFe Group 4*** | 73 | 99 | 92 |
| ***Fe*** | 0 | 0 | 0 |

**Supplementary Figure 1** Map of the Eastern Gulf of Mexico (GoM) showing the studied three sampling locations (E26, E29 and E44) and bathymetry of the study area.

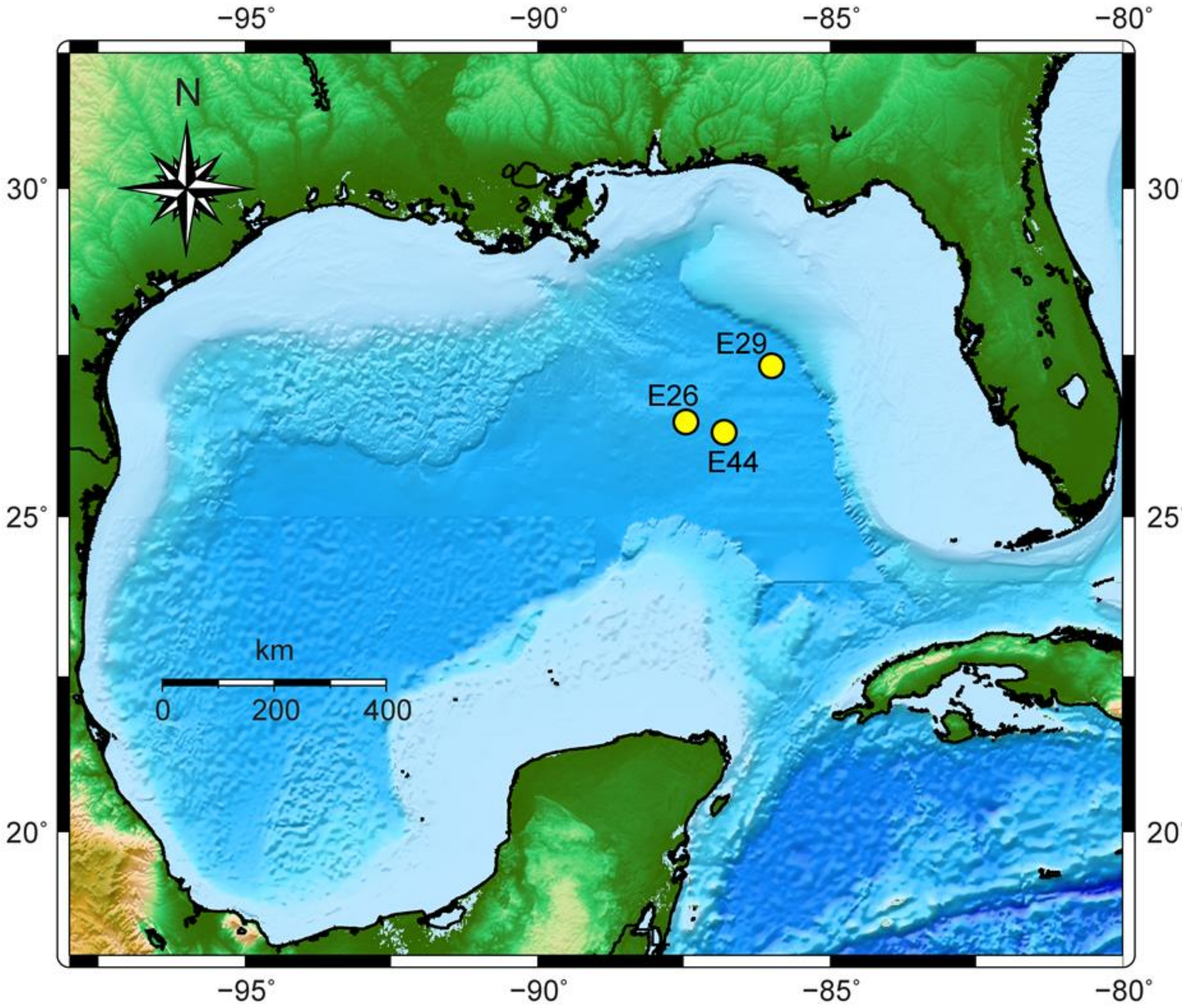

**Supplementary Figure 2** Phylum-level community composition of classified bacterial (left) and archaeal (right) 16S rRNA gene reads from separate bacterial and archaeal amplicon libraries. Detailed numbers can be found in Supplementary Table 3.

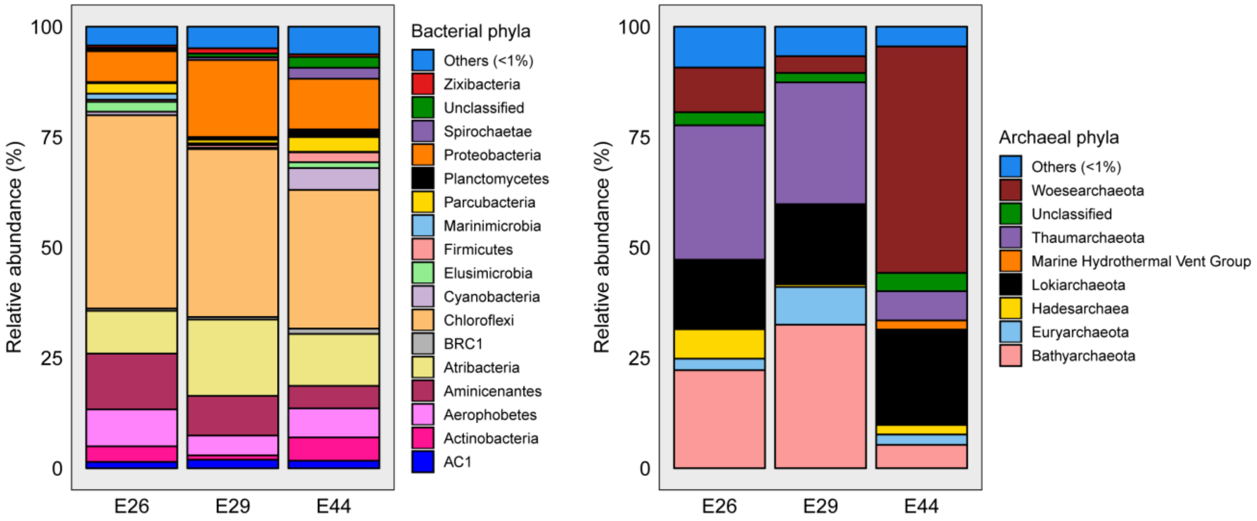

**Supplementary Figure 3** Phylogenetic relationship of putative glycyl-radical enzymes in MAGs with those of akyl-/arylalkylsuccinate synthases. Reference sequences show canonical alkane succinate synthase (AssA), 4-hydroxylbenzyl succinate synthase (HbsA), benzyl succinate synthase (BssA), and homologous putative alkane-degrading enzymes from *Vallitalea guaymasensis* L81 and *Archaeoglobus fulgidus* VC-16. The tree was constructed with the maximum-likelihood method (Poisson correction model) and is bootstrapped with 50 replicates. Sequences of pyruvate formate lyase (Pfl) from *E. coli* were used as an outgroup (not shown).

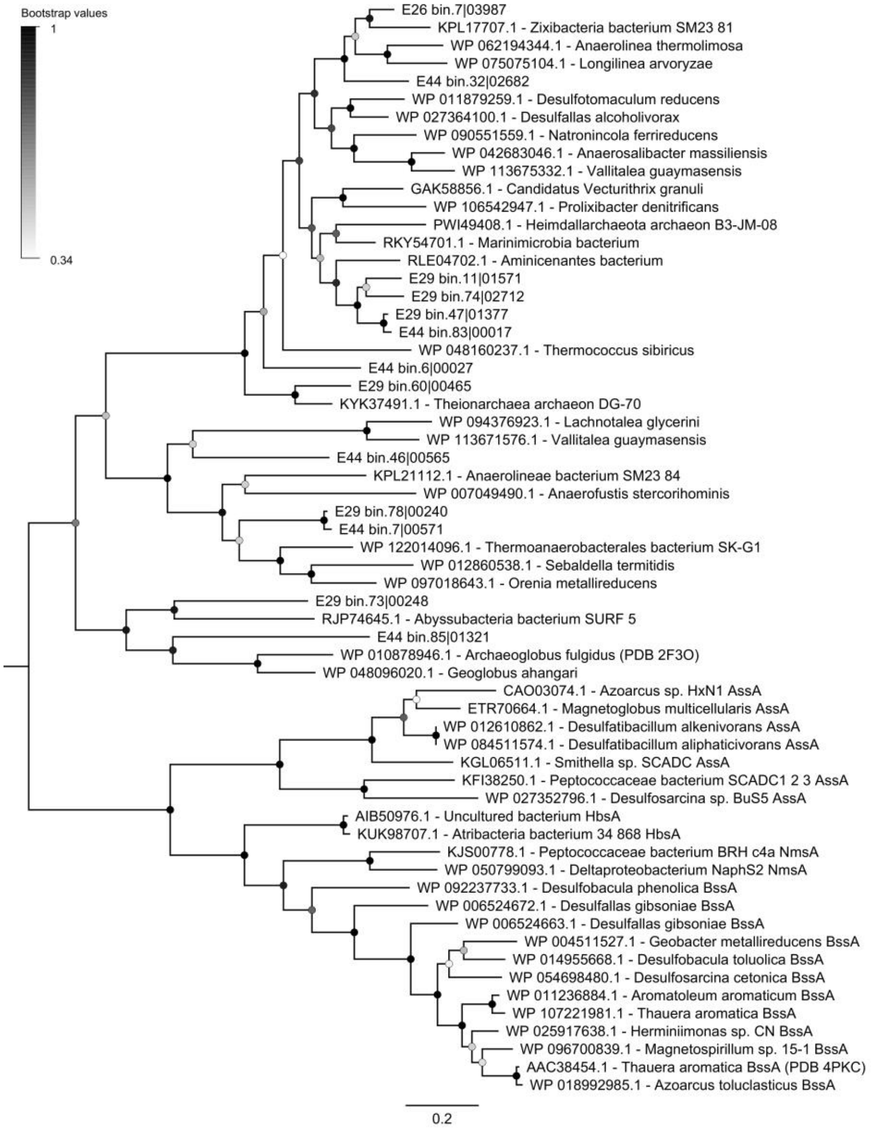

**Supplementary Figure 4** Phylogenetic relationship of identified genes in MAGs with currently-known molybdenum cofactor-containing hydrocarbon dehydrogenases. Reference sequences show the catalytic subunits of characterized cymene dehydrogenase (CmdA), alkane $C_2$-methylene hydroxylase (AhyA), and ethylbenzene dehydrogenase (EbdA/EbdA2) enzymes from other hydrocarbon-degrading bacteria. The tree was constructed with the maximum-likelihood method (Poisson correction model) and is bootstrapped with 50 replicates. Sequences of perchlorate reductase (PcrA) from *Dechloromonas aromatica* RCB were used as an outgroup (not shown).

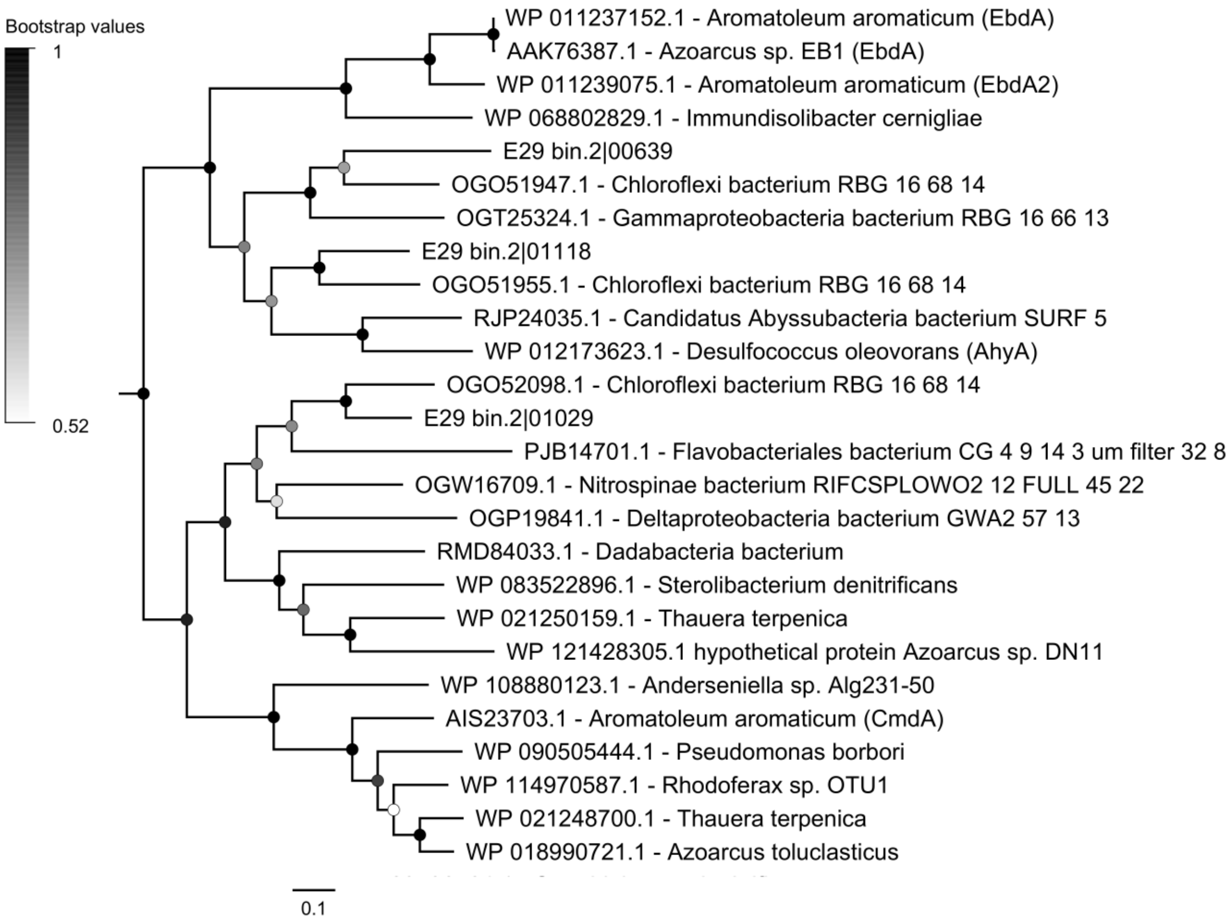

**Supplementary Figure 5** Protein presence/absence matrix for benzoyl-CoA anaerobic biodegradation pathway. The MAGs were shown only if it was at least partially complete (presence of at least three subunits within one cluster for BcrABCD). Presence of genes isindicated by blue boxes. Gene names: Bcr, benzoyl-CoA reductase; Oah, 6-oxo-cyclohex-1-ene-carbonyl-CoA hydrolase; Dch, cyclohex-1,5-diencarbonyl-CoA hydratase; Had, 6-hydroxycyclohex-1-ene-1-carbonyl-CoA dehydrogenases. More details about these functional genes and pathways can be found in the text and in Supplementary Table 6.

| **Bin No.** | **Lineage** | **Bcr** | **Dch** | **Had** | **Oah** |
|---|---|---|---|---|---|
| E29_bin47 | *Aminicenantes* | ■ | ■ | ■ | ■ |
| E26_bin7 | *Anaerolineales* | ■ | ■ | ■ | ■ |
| E29_bin75 | *Dehalococcoidia* | ■ | □ | ■ | □ |
| E44_bin56 | *Dehalococcoidia* | ■ | ■ | □ | □ |
| E44_bin89 | *Dehalococcoidia* | ■ | □ | ■ | □ |
| E44_bin91 | *Desulfobacteraceae* | ■ | ■ | □ | □ |
| E29_bin36 | *TA06* | ■ | ■ | ■ | □ |
| E44_bin18 | *TA06* | ■ | ■ | ■ | ■ |
| E26_bin22 | *Bathyarchaeota* | ■ | ■ | ■ | ■ |
| E29_bin60 | *Bathyarchaeota* | ■ | ■ | ■ | ■ |
| E44_bin43 | *Bathyarchaeota* | ■ | ■ | ■ | ■ |
| E29_bin30 | *Thermoplasmata* | ■ | □ | ■ | ■ |

**Supplementary Figure 6** Protein presence/absence matrix for reductive acetyl-CoA (Wood-Ljungdahl) pathway. Presence of genes is indicated by blue boxes. Columns correspond to the following enzymes: 1, formate dehydrogenase (Fhd) / formylmethanofuran dehydrogenase (Fwd); 2, formate-tetrahydrofolate synthetase (Fhs) / formylmethanofuran:tetrahydromethanopterin formyltransferase (Ftr); 3, methylene-tetrahydrofolate dehydrogenase (FolD) / N5,N10-methenyltetrahydromethanopterin cyclohydrolase (Mch); 4, methylene-tetrahydrofolate dehydrogenase (FolD) / methylenetetrahydromethanopterin dehydrogenase (Mtd); 5, methylenetetrahydrofolate reductase (Met) / N5,N10-methylenetetrahydromethanopterin reductase (Mer), 6, acetyl-CoA synthetase (Acs), 7 carbon monoxide dehydrogenase (Cdh).

| Bin No. | Lineage | Fhd Fwd | Fhs Ftr | FolD Mch | FolD Mtd | Met Mer | Acs | Cdh |
|---|---|---|---|---|---|---|---|---|
| E29_bin7 | *Actinobacteria* | | | | | | | |
| E29_bin77 | *Actinobacteria* | | | | | | | |
| E44_bin5 | *Actinobacteria* | | | | | | | |
| E29_bin28 | *Aerophobetes* | | | | | | | |
| E29_bin52 | *Aerophobetes* | | | | | | | |
| E29_bin78 | *Aerophobetes* | | | | | | | |
| E44_bin3 | *Aerophobetes* | | | | | | | |
| E44_bin92 | *Aerophobetes* | | | | | | | |
| E29_bin47 | *Aminicenantes* | | | | | | | |
| E29_bin74 | *Aminicenantes* | | | | | | | |
| E44_bin65 | *Atribacteria* | | | | | | | |
| E26_bin7 | *Anaerolineales* | | | | | | | |
| E29_bin16 | *Anaerolineales* | | | | | | | |
| E29_bin24 | *Anaerolineales* | | | | | | | |
| E44_bin32 | *Anaerolineales* | | | | | | | |
| E44_bin81 | *Anaerolineales* | | | | | | | |
| E25_bin16 | *Dehalococcoidia* | | | | | | | |
| E29_bin15 | *Dehalococcoidia* | | | | | | | |
| E29_bin2 | *Dehalococcoidia* | | | | | | | |
| E29_bin42 | *Dehalococcoidia* | | | | | | | |
| E29_bin44 | *Dehalococcoidia* | | | | | | | |
| E29_bin54 | *Dehalococcoidia* | | | | | | | |
| E29_bin73 | *Dehalococcoidia* | | | | | | | |
| E29_bin75 | *Dehalococcoidia* | | | | | | | |
| E44_bin26 | *Dehalococcoidia* | | | | | | | |
| E44_bin27 | *Dehalococcoidia* | | | | | | | |
| E44_bin29 | *Dehalococcoidia* | | | | | | | |
| E44_bin46 | *Dehalococcoidia* | | | | | | | |
| E44_bin56 | *Dehalococcoidia* | | | | | | | |
| E44_bin88 | *Dehalococcoidia* | | | | | | | |
| E44_bin89 | *Dehalococcoidia* | | | | | | | |
| E29_bin43 | *Cloacimonetes* | | | | | | | |
| E44_bin80 | *Cloacimonetes* | | | | | | | |
| E29_bin65 | *Desulfobacteraceae* | | | | | | | |
| E44_bin91 | *Desulfobacteraceae* | | | | | | | |
| E44_bin39 | *Planctomycetes* | | | | | | | |
| E29_bin36 | *TA06* | | | | | | | |
| E44_bin18 | *TA06* | | | | | | | |
| E44_bin34 | *Heimdallarchaeota* | | | | | | | |
| E29_bin63 | *Lokiarchaeota* | | | | | | | |
| E44_bin85 | *Lokiarchaeota* | | | | | | | |
| E44_bin77 | *Thorarchaeota* | | | | | | | |
| E26_bin22 | *Bathyarchaeota* | | | | | | | |
| E26_bin4 | *Bathyarchaeota* | | | | | | | |
| E29_bin39 | *Bathyarchaeota* | | | | | | | |
| E29_bin53 | *Bathyarchaeota* | | | | | | | |
| E29_bin60 | *Bathyarchaeota* | | | | | | | |
| E44_bin4 | *Bathyarchaeota* | | | | | | | |
| E44_bin43 | *Bathyarchaeota* | | | | | | | |
| E29_bin30 | *Thermoplasmata* | | | | | | | |
